# Direct RNA-RNA interaction between Neat1 and RNA targets, as a mechanism for RNAs paraspeckle retention

**DOI:** 10.1101/2020.10.26.354712

**Authors:** Audrey Jacq, Denis Becquet, Séverine Guillen, Bénédicte Boyer, Maria-Montserrat Bello-Goutierrez, Jean-Louis Franc, Anne-Marie François-Bellan

## Abstract

Paraspeckles are nuclear ribonucleic complex formed of a long non-coding RNA, nuclear-enriched abundant transcript one (Neat1) and associated RNA-binding proteins (RBP) whose cellular known functions are to sequester in the nucleus both proteins and RNAs. However, how RNAs are bound to paraspeckles is largely unknown. It is highly likely that binding of RNAs may occur via interactions with RBPs and accordingly, two structures present in the 3’UTR of some RNAs have been shown to allow their association to paraspeckles via protein binding. However, Neat1 could also be involved in the targeting of RNAs through direct RNA-RNA interactions. Using a RNA pull-down procedure adapted to select only RNAs engaged in direct RNA-RNA interactions and followed by RNA-seq we showed that in a rat pituitary cell line, GH4C1 cells, 1791 RNAs were associated with paraspeckles by direct interaction with Neat1. Neat1 was actually found able to bind more than 30% of the total transcripts targeted by the paraspeckles, we have identified in this cell line in a previous study. Furthermore, given the biological processes in which direct RNAs targets of Neat1 were involved as determined by gene ontology analysis, it was proposed that Neat1 played a major role in paraspeckle functions such as circadian rhythms, mRNA processing, RNA splicing and regulation of cell cycle. Finally, we provided evidence that direct RNA targets of Neat1 were preferentially bound to the 5’ end of Neat1 demonstrating that they are located in the shell region of paraspeckles.

## INTRODUCTION

Paraspeckles are nuclear ribonucleoprotein complexes found in almost all cell lines and tissues from mammals, except for embryonic stem cells (for review see 1, 2). These nuclear bodies are usually detected as a variable number of discrete dots found in close proximity to nuclear speckles (3, 4). The structural element of paraspeckles is a long noncoding RNA, nuclear-enriched abundant transcript one (Neat1). The locus of Neat1 generates two major isoforms, the short one Neat1-1 (previously named MENε) and the long one Neat1-2 (MENβ) which are transcribed from the same promoter (5, 6). It is known that paraspeckle proteins (PSPs) accumulate on Neat1-2 isoform to form the paraspeckles (7) and that not less than 60 proteins have been identified so far in these nuclear bodies (8). Among them two RNA-binding proteins (RBP), HNRNPK and RBM14, as well as two proteins of the Drosophila Behavior Human Splicing (DBHS) family, NONO and SFPQ, have been shown to be essential for the formation and maintain of paraspeckles (9). These well organized structures can be subdivided into two zones, the core and the shell, the later containing both the 5’ end of Neat1-1 and the 5’ and 3’ ends of Neat1-2 (10, 11).

In addition to proteins, paraspeckles have been shown to retain RNAs in the nucleus. In a rat pituitary cell line, GH4C1 cells, we have previously shown that the expression of both Neat1 and four major PSPs followed a circadian pattern that leads to rhythmic variations in paraspeckle number within the cells (12). As a consequence, paraspeckles rhythmically retain target RNAs in the nucleus of the cells, leading to the rhythmic expression of the corresponding genes (13). However, how target mRNAs are bound to paraspeckles is largely unknown. The presence of duplex structures in the target RNAs (14) is a feature that can lead to paraspeckle retention, as it was shown to be the case for the mouse cationic amino acid transporter 2 transcribed nuclear-RNA (Ctn-RNA) and the human RNAs Nicolin 1 (NICN1) and Lin28 (15, 16). Indeed these later RNAs contain a dsRNA structure resulting from inverted repeated short interspersed nuclear elements (IRSINEs) in their 3’-UTR (15). In human cells, hundreds of genes contain inverted repeated IRSINEs (mainly IRAlu elements) in their 3’-UTRs (16). However, unexpectedly we didn’t find IRSINEs in 3’-UTR of the 4268 RNAs, we previously identified as paraspeckle RNAs targets using a Neat1 RNA pull-down procedure (17) followed by RNA-sequencing in GH4C1 cells (12). By contrast, a sequence motive of 15 nucleotides identified in the 3’UTR of more than 30% of the 4268 RNAs that are paraspeckle targets may be involved in the nuclear retention of RNAs by paraspeckles through its binding by PSP component HNRNPK (18). However, whereas PSPs probably play a crucial role in binding RNA targets, it is also possible that the lncRNA Neat1, in addition to its structural role, may also be directly involved in the binding by base pairing of RNAs and consequently in their retention in paraspeckles.

To test this hypothesis, we adapted the Neat1 RNA pull-down procedure we previously described (17) by treating GH4C1 cells with psoralen, which only fix the RNA-RNA interactions by crosslinking the uracil-uracil bounds (19). We found 1791 RNAs directly bound by Neat1, which represent more than 30% of the total paraspeckle targets previously identified (12). By gene ontology analysis, the direct RNAs targets of Neat1 were further shown involved in major paraspeckle functions such as circadian rhythms, mRNA processing, RNA splicing and regulation of cell cycle, underlying the crucial role Neat1 played in these functions by means of RNA-RNA interactions. Since direct RNA targets of Neat1 were shown preferentially bound to the 5’ end of Neat1, it is proposed that these direct RNA targets were mainly localized in the shell region of paraspeckles.

## MATERIALS AND METHODS

### Cell line culture

GH4C1 cells, a rat pituitary somatolactotroph line, were obtained from ATCC (CCL-82.2, lot number: 58945448) with certificate analysis and were confirmed to be free of mycoplasma (MycoAlert). They were grown in 10 cm cell dishes, in an incubator at 37°C, saturated with H_2_O and with 5% CO_2_. The HamF10 cell medium was supplemented with 15% horse serum, 2% fetal calf serum, 0.5% streptomycin and 0.5% penicillin. To synchronize cells between themselves and to be able to select the time of maximum Neat1 expression (12), GH4C1 cells were transferred to fresh medium 24 to 30 h before crosslinking.

### Neat1 pull down

LncRNA pull-down (17) is a hybridization-based strategy that uses complementary oligonucleotides to purify lncRNA together with its targets from reversibly cross-linked extracts. In cross-linked extracts, it is expected that some regions of the RNA will be more accessible for hybridization than others due in particular to secondary structure. To design oligonucleotides that target these regions and then can hybridize specifically to lncRNA Neat1, we modeled the secondary structure of Neat1 RNA using the RNAstructure software (20). Two pools (A and B) of 6 antisens DNA oligonucleotide probes that target accessible regions throughout the length of the lncRNA Neat1 were designed and used for the specific Neat1 RNA pull-down (Figure 1; Supplemental Table 1). All these probes were biotinylated at the 3’ end (IDT, Coralville, Iowa, USA)

**Figure 1.**
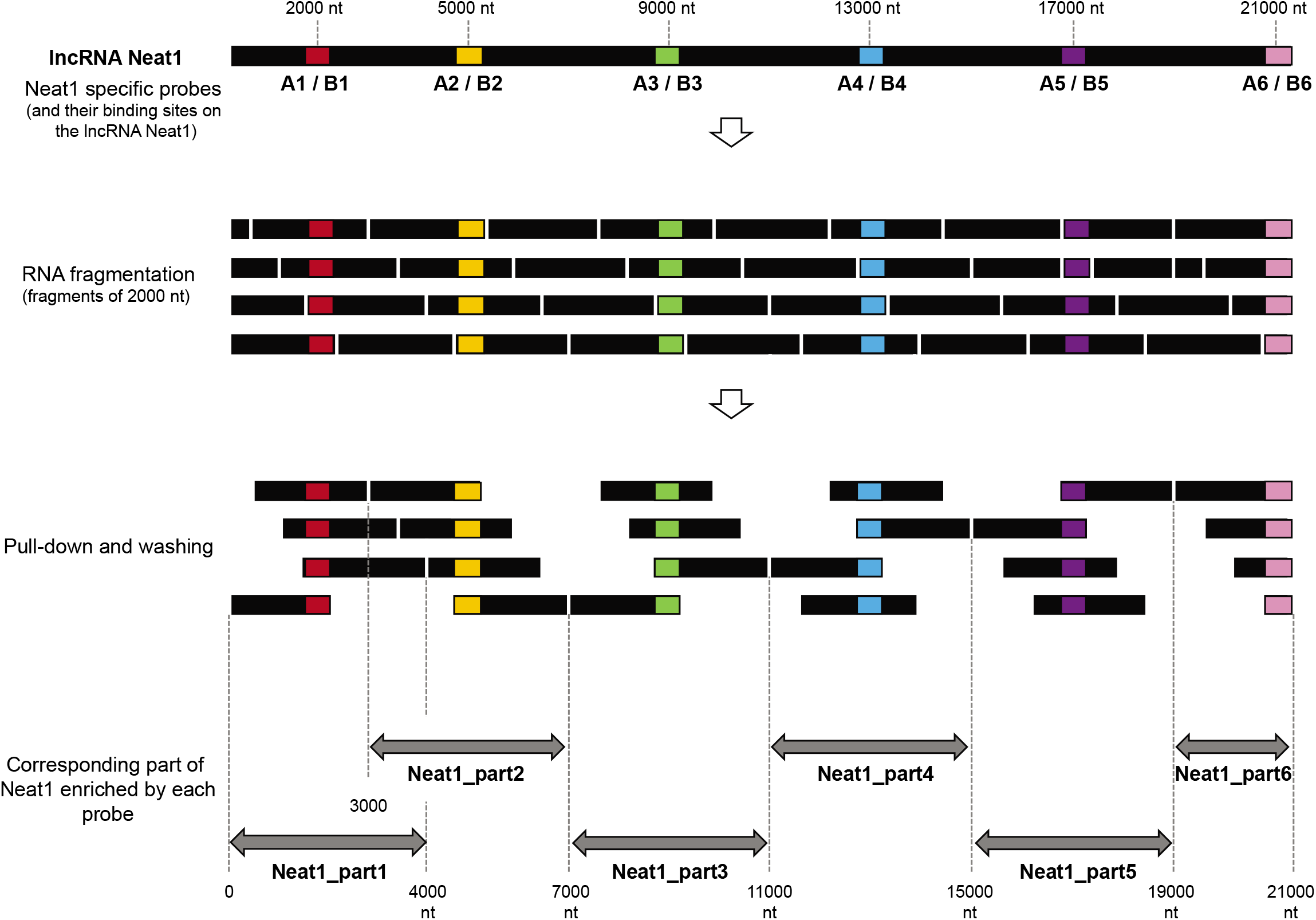
Schematic representation of the pull-down of 6 parts of Neat1 by 6 couples of specific probes. Schematic localization of the binding sites for 6 couples of specific antisens oligonucleotides designed along the length of Neat1. After Neat1 was fragmented in around 2000 nucleotides fragments following the sonication procedure, the different fragments as well as the corresponding parts of Neat1 pulled-down by the different couples of probes (A and B) are shown.

Twenty-four hours minimum after the fresh medium replacement, GH4C1 cells were rinsed one time with cold PBS with Ca^++^ and Mg^++^, then incubate with 0.1 mg/ml psoralen-derived molecule (4’-Aminomethyltrioxsalen hydrochloride, Sigma) and placed 30 min in the dark, in an incubator at 37 °C with 5% CO_2_. Then cells were placed on ice at 2.5 cm of the 365 nm UV tubes in an UV Stratalinker 1800 (Stratagene, San Diego, CA, USA) at 5000 μJ x 100, two times 10 min. Cross-linked cells were rinsed with PBS, scraped and separated by centrifugation (400 g for 5 min at 4 °C). Cell pellets were stored at −80°C.

To prepare lysates, cells pellets were suspended in Proteinase K buffer (100 mM NaCl, 10 mM Tris-HCl, pH 7.0, 1 mM EDTA, 0.5% SDS and 5 μl/ml RNase-Out). 0.1 μg/μl of Proteinase K (Ambion, Waltham, MA, USA) were added and the lysates were incubated 45 min at 50 °C following by 13 min at 95 °C, to inactivate the Proteinase K. Lysates were separated into different sample tubes and supplemented with 2 volumes of Hybridization buffer (700 mM NaCl, 70 mM Tris-HCl, pH 7.0, 1 mM EDTA, 1.25% SDS, 5 μl/ml RNase-Out and 15% formamide). Diluted lysates were sonicated using BioruptorPlus (Diagenode, Seraing, Belgium) by 2 pulses of 30 seconds allowing complete lysate solubilization. The size of the RNA fragments generated was verified on a gel and corresponded to about 2000 nucleotides. 20 μl of diluted samples were collected and stored at −80°C to serve as the input control samples. Specific probes (pool A of 6 specific probes and pool B of 6 specific probes, 90 pmol in total) or non-specific probe (90 pmol) were added to the diluted lysates, which were mixed by end-to-end rotation at room temperature for 4 h. After one washing with the Hybridization buffer, Streptavidin-magnetic C1 beads (Dynabeads MyOne Streptavidin C1, Thermo Fisher Scientific) were re-suspended in Hybridization buffer and added to hybridization reaction (40 μl per 100 pmol of probes). The whole reaction was incubated under agitation overnight at room temperature. Beads-biotin-probes-RNA adducts were captured by magnets (Promega, Madison, WI, USA) and washed five times with the Wash buffer (2X SSC, 0.5% SDS). After the last wash, buffer was removed carefully. For RNA elution, beads and input samples were suspended in Proteinase K buffer with 1 μg/μl proteinase K. After incubation at 45 °C for 45 min followed by 10 min at 95 °C, RNA was isolated using NucleoSpinRNA XS kit (Macherey-Nagel, Dueren, Germany). Eluted RNA was subject to RT-qPCR or RNA sequencing for the detection of enriched transcripts.

### RNA expression analysis by RT-qPCR

Total RNA was used for cDNA synthesis performed with a High Capacity RNA to cDNA kit (Applied Biosystem, Waltham, MA, USA). Real-time PCR was performed on a CFX96^TM^ Real-Time PCR system (Bio-Rad, Hercules, CA, USA) using iTaq^TM^ Universal SYBR^®^ Green Supermix (Bio-Rad). The sequences of the primers used in qPCR are given in Supplemental Table 1. mRNA accumulation was normalized to mRNA levels in inputs and/or non-specific probe sample.

### RNA expression analysis by RNA sequencing

The construction of Illumina DNA libraries and the sequencing from RNA pools obtained in triplicate for each of the two pools of specific oligonucleotides were performed by Genewiz (Leipzig, Germany). RNA recovery after use of the non-specific probe was too low to allow the construction of a library. Libraries were prepared with Illumina Sample Preparation kit with rRNA depletion. Strand-specific RNA-seq was done on Illumina HiSeq 2500, with a read length of 2×150 bp (30 million reads per sample on average were obtained).

Analyses were performed on a local instance of Galaxy. After quality control checks by FastQC and check for adapter content with Trimmomatic, paired-end reads were aligned to the Rat reference genome (Rnor_6.0.80, Ensembl) using Star (21). Then, FeatureCounts (22) was used to quantify the number of counts for each gene. In order to assess the specificity of the Neat1 RNA pull-down we crossed the results obtain with the two pools of oligonucleotides to generate the list of transcripts associated to the lncRNA Neat1.

The RNA sequencing data are available at Gene Expression Omnibus (GEO) (accession number n°GSE160069).

### Statistical analysis

For data obtained by qPCR analysis, significant differences between groups were determined using one-way or two-way ANOVA as needed (Prism 6 software). Values were considered significantly different for p-value < 0.05(*).

## RESULTS

### Identification of RNAs engaged in RNA-RNA interactions with Neat1

To identify RNAs that were bound to paraspeckles through direct interaction with Neat1 itself, RNA pull-down experiments were performed according to a protocol we have previously described (17) except that to fix the cells, psoralen, a molecule that crosslinks only RNA-RNA interactions (19) was used instead of paraformaldehyde (PFA). After crosslinking, cells were lysed, and extracts obtained were submitted to sonication in order to obtain RNA fragments of about 2000 nucleotides. Probably because of the extended length of Neat1-2 (21 kb), this step proved to be indispensable for its efficient pull-down. Because of the fragmentation of Neat1-2 generated by the sonication procedure, it may be assumed that a specific antisens probe could pull-down fragments of Neat1 that were located up to 2000 nt upstream and 2000 nt downstream from the site of probe binding (Figure 1). Consequently, a specific probe could pull-down fragments corresponding to a Neat1 part of 4000 nt length. In order to ensure the pull-down of the entire length of Neat1-2 transcript, we designed 12 Neat1-specific biotinylated probes able to bind 6 different Neat1 parts (a couple of A and B probes for each part) (Supplemental Table1). The parts of Neat1-2 that were pulled-down by each couple of probes were delineated in Figure 1.

By qPCR using specific primers that target each of the 6 parts of Neat1 (Supplemental Table1), we verified that each couple of probes (A-B) was able to specifically bind the part of Neat1 against which it was designed as illustrated in RNA pull-down performed with each of the six A probes (Figure 2).

**Figure 2.**
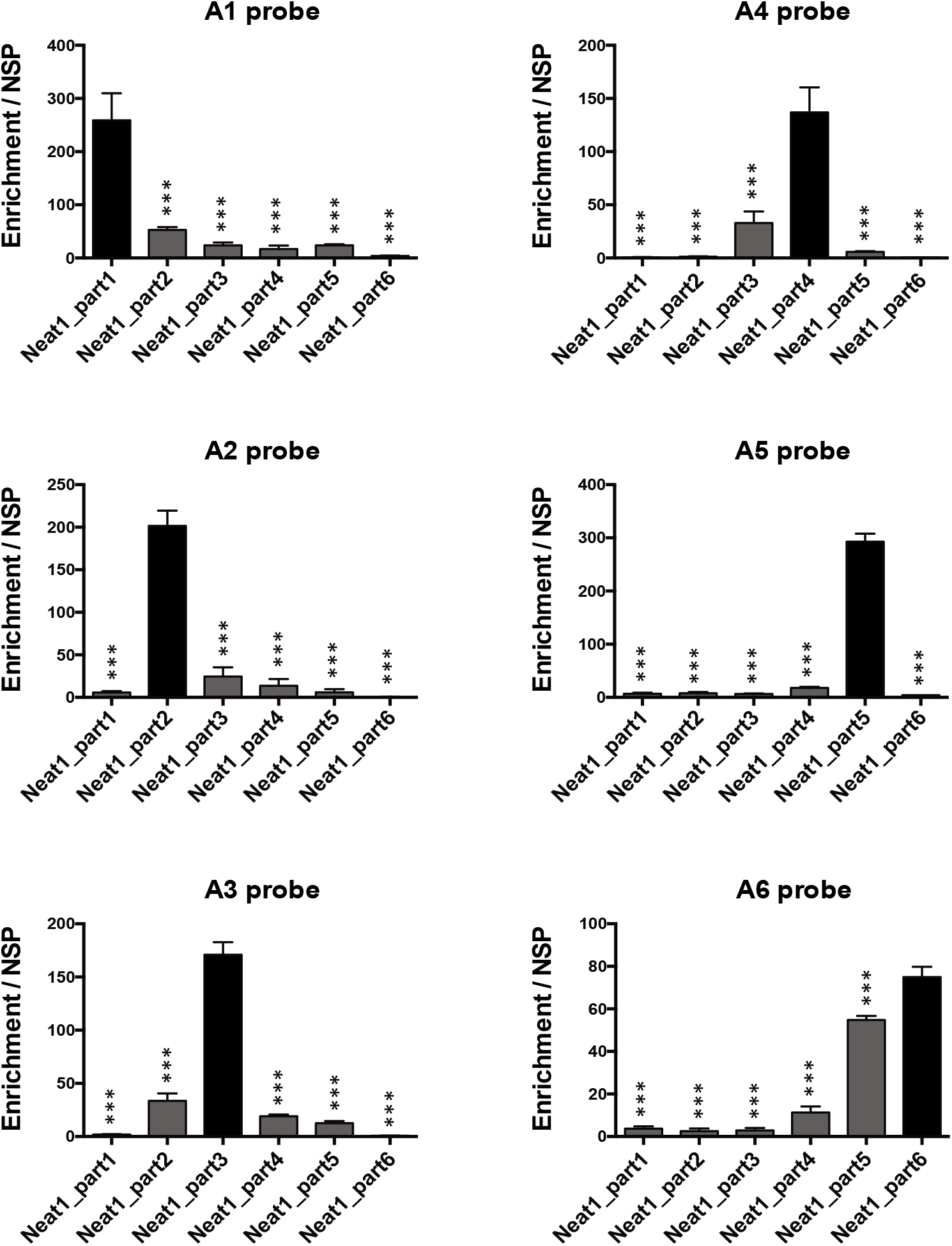
Pull-down of 6 parts of Neat1 performed with each of the six A probes. Specific parts of Neat1 pulled down by each specific probe from the set of A probes. The specific enrichment of the different parts of Neat1 was determined by RT-qPCR after pulldown with each specific probe compared to a non-specific probe (NSP). ***p<0.001.

With the aim to recover all RNAs that were directly bound to Neat1 throughout its length, all six A probes on the one hand and all six B probes on the other hand were pooled and used as pool A and pool B in Neat1 pull-down experiments. The efficacy of these two pools of probes to pull-down Neat1 was verified (Supplemental Figure 1). Every 6 parts of Neat1 were shown to be efficiently pulled-down with the pool A and the pool B of probes even if depending on the part of Neat1 considered, the efficacy could differ between pool A and pool B (Supplemental Figure 1). To assess the specificity of the Neat1 RNA pull-down experiments performed, results obtained with these two different pools were crossed. Experiments using a non-specific probe were also done and gave rise to a very low RNA recovery as compared to the two specific probe sets. Accordingly, only the RNAs pulled-down with the two A and B pools were analyzed by RNA-seq.

After the creation and sequencing of the libraries, the reads were aligned on the rat genome using Star (21) and FeatureCounts (22) was used to quantify counting reads as a measure of RNA precipitated (Supplemental Table 2). Only transcripts exhibiting a substantial number of counts (>200) were selected. After use of the pool A of probes and the pool B of probes, a List-A composed of 2564 RNAs and a List-B composed of 2381 RNAs were established, respectively (Supplemental Table 2). The specificity of this Neat1 RNA pull-down experiment was assessed by crossing the two lists obtained. 1791 RNAs were common to both lists and named List-AB. This represented 70% of List-A and 75% of List-B (Figure 3A – Supplemental Table 2). To validate the results obtained after RNA-seq, we selected 13 RNAs that were analyzed by RT-qPCR after Neat1 RNA pull-down experiments performed with the pool A of specific probes or a non-specific probe. Among these 13 RNAs, 10 were included in the List-AB, and 3 (Prkcb, Rpa1 and Tapbp) weren’t (neither in List-A nor in List-B). The 10 RNAs from the List-AB which were sorted here according to the number of counts obtained in RNA-seq, from largest to smallest, exhibited an enrichment with respect to the non-specific probe that was significantly higher compared to that of the 3 negative control RNAs that were not included in the List-AB (F_1/76_ = 11.06, p=0.0014); magnitude of the enrichment however was shown to greatly differ between the 10 RNAs and to be independent on the number of counts obtained in RNA-Seq (Figure 3B). By selecting 6 RNAs from the List-AB according to their RNA counts found in RNA-seq analysis (2 RNAs with 3000, 1300 or 400 counts, respectively), we further showed by RT-qPCR that a same number of counts in RNA-seq could be associated with very different enrichment scores and RNAs with very different number of counts could exhibit a same enrichment score (Supplemental Figure 2). It then appeared that the enrichment score of RNAs was not relative to their level of expression.

**Figure 3.**
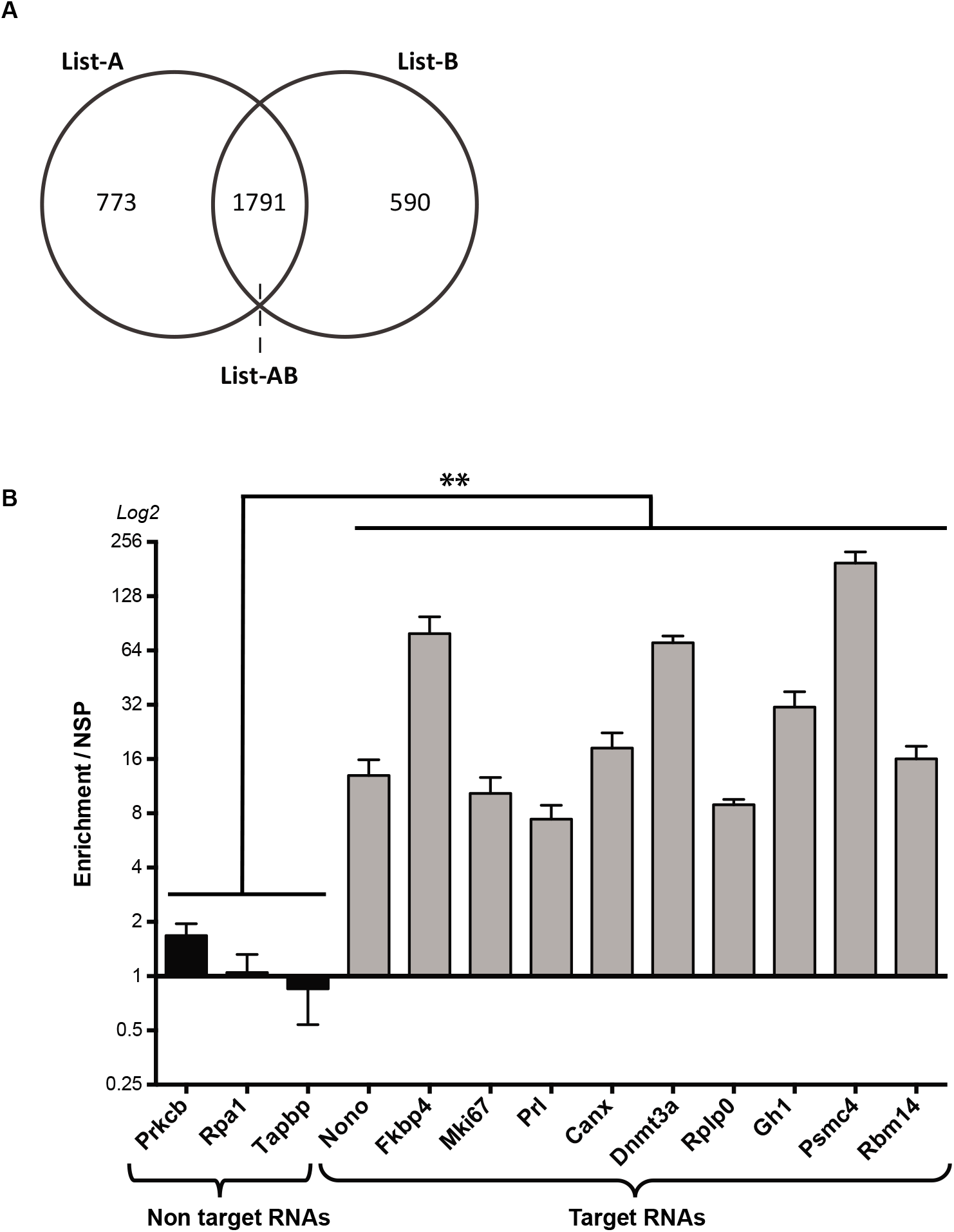
RNAs directly bound by the lncRNA Neat1. **A.** Venn diagram illustrating the overlap between the lists of transcripts targeted by pool A (List-A) and pool B (List-B) of Neat1 specific probes, respectively. The list of RNAs commons to List-A and to List-B is refereed to List-AB. **B.** Validation by RT-qPCR of results from RNA-seq. After selection of 13 RNAs [10 RNAs targeted by Neat1 (found in List-AB and sorted according to the number of counts obtained in RNA-seq from largest to smallest) and 3 non target RNAs (found neither in List-A nor in List-B)], the enrichment obtained after Neat1 RNA pull-down with the pool A of specific probes relative to a non-specific probe is shown to be significantly different in Neat1 targets compared to non-targets (F_1,76_=11.06 p=0.0014). **p<0.01.

### Contribution of direct Neat1 RNA targets to the total RNAs targets of paraspeckles

To evaluate the proportion of paraspeckle RNA targets in GH4C1 cells that were directly connected by Neat1 through RNA-RNA interactions, we compared our present list of direct Neat1 targets with the dataset of total paraspeckle RNA targets we have previously established using Neat1 RNA pull-down experiments performed on cells fixed by PFA (12).

Indeed, while psoralen used to fix the cells in the present study allowed to selectively crosslink only RNA-RNA interactions, PFA permitted to crosslink both RNA-RNA and RNA-protein interactions leading to the pull-down of both direct and indirect Neat1 targets. By crossing over the list of 1791 RNAs directly targeted by Neat1 (List-AB) with the list of 4268 RNAs previously identified as paraspeckle targets (12), 1398 genes appeared common to the two lists (Figure 4A; Supplemental Table 3). This represented almost 33% of the paraspeckle targets. However, while it may be noticed that 78% of the List-AB was found to be included in our previous list of RNA paraspeckle targets, 22% (393 RNAs) of the direct targets of Neat1, as reported here, were previously unidentified targets of paraspeckles.

### Functional analysis of direct Neat1 RNA targets

Enrichment in specific biological functions and pathways of the 1791 RNAs from the List-AB was analyzed using the DAVID Bioinformatics Resources (23). DAVID provides a clustering function that forms sets of overlapping gene categories. Circadian rhythms, mRNA processing, RNA splicing and regulation of cell cycle were found among the most prominent enriched annotation clusters in biological processes (p<0.05; Figure 4B – Supplemental Table 4). KEGG pathways classification showed that circadian entrainment, cell cycle, microRNA in cancer, RNA transport and spliceosome were significantly enriched in our data set (p<0.05; Figure 4C – Supplemental Table 4).

### Mapping of Neat1 regions involved in RNA-RNA interactions

To assess whether peculiar regions of Neat1 were involved in RNA-RNA binding, a single-probe RNA pull-down experiment was performed using the 6 probes of pool A separately (from A1 to A6). We selected 5 different RNAs from the List-AB (Canx, Fkbp4, Nono, Prl, Rbm14), Malat1 which is not officially annotated in the rat genome but was previously shown to be associated with paraspeckle (12) and Rpa1 which was not included in the List-AB. Enrichment analysis was performed by qPCR as compared to the non-specific probe (Figure 5). Each value of enrichment was normalized to the percentage of recovery of the corresponding Neat1 part to ensure that results did not depend on the degree of recovery of each part of Neat1 by its corresponding probe. The results showed that the 5 genes of the List-AB as well as Malat1 bound to the Neat1 5’ end (Neat1_part1 bound by probe A1). In a more discrete way, it could be noticed that the 3’ end of Neat1-2 (Neat1_part5 and Neat1_part6 targeted by probes A5 and A6, respectively) seemed also able to bind some of the 6 RNAs although enrichments didn’t reach statistical significance (Figure 5). By contrast Rpa1 was never targeted by any of the 6 probes (Figure 5).

**Figure 4.**
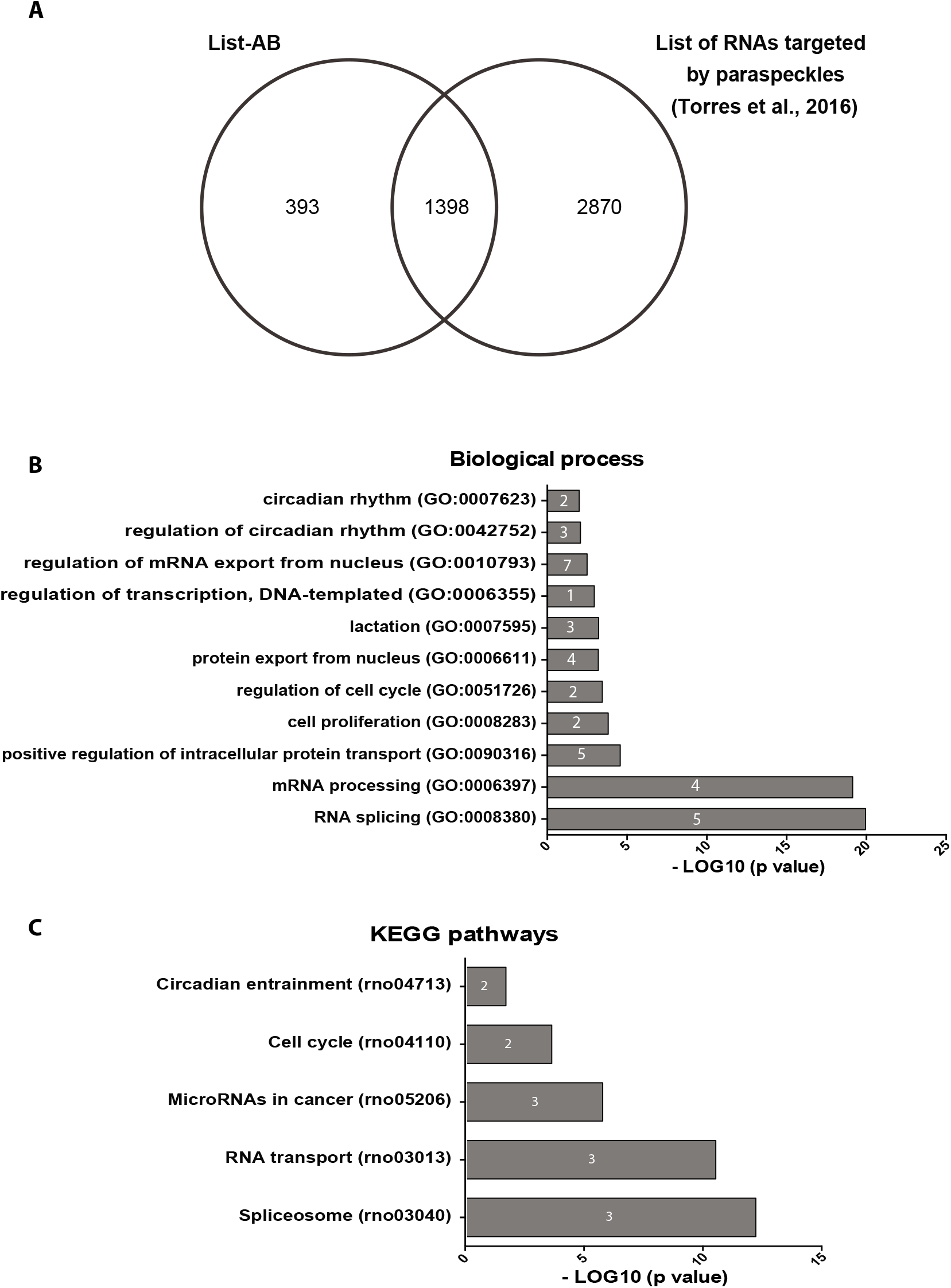
Functional analysis of direct Neat1 RNA targets. **A.** Venn diagram representation of the overlap between total RNA paraspeckle targets (12) and direct Neat1 RNA targets (List-AB). **B-C** David analysis of direct Neat1 RNA targets. **B.** Examples of biological process terms identified by the Gene Ontology analysis and sorted according to their p-value in Log10; fold enrichment is given inside histogram bars. **C.** Pathways identified by the KEGG Classification System and sorted according to their p-value in Log10; fold enrichment is given inside histogram bars.

**Figure 5.**
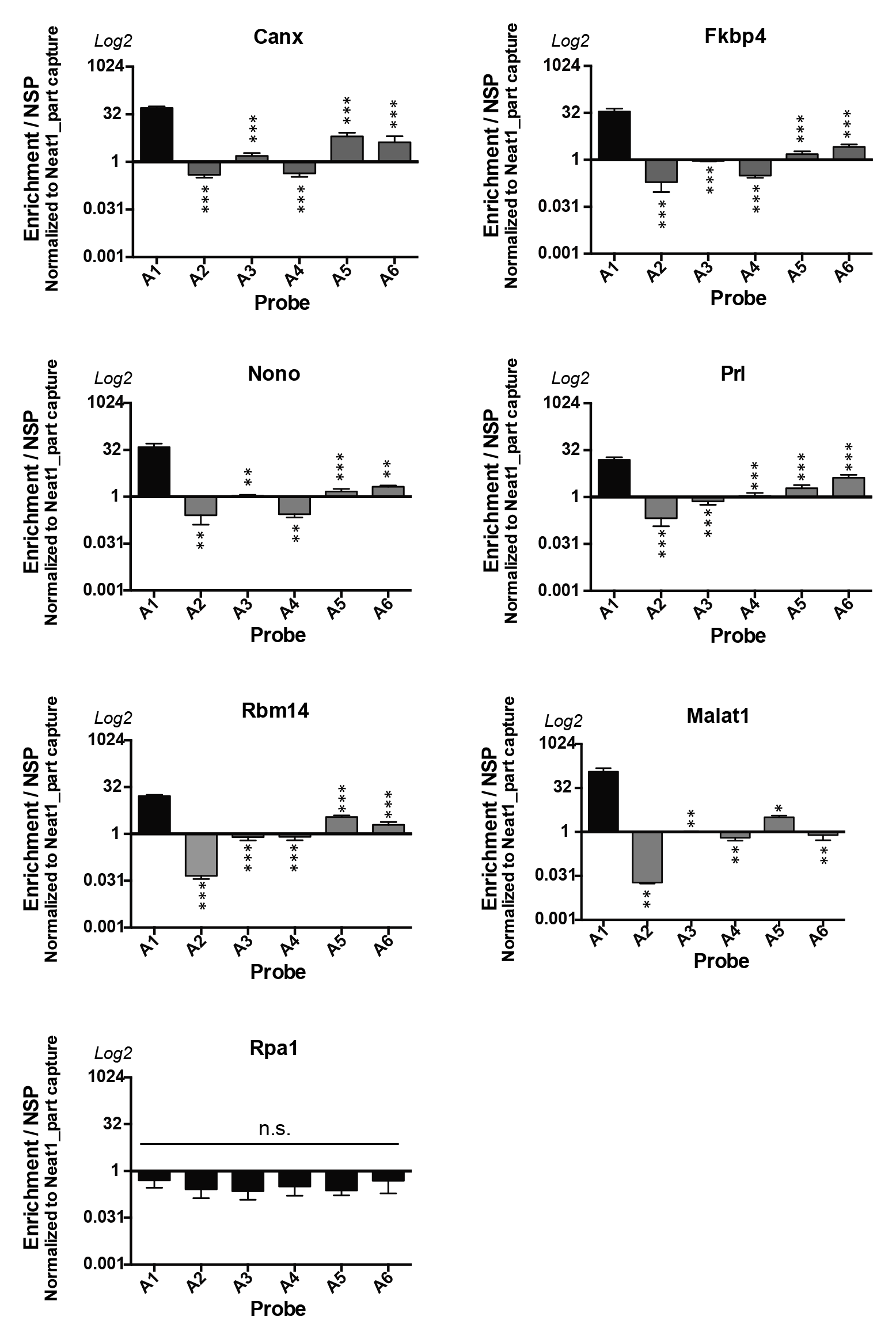
Involvement of the 5’ region of Neat1 in direct RNA interactions. Each Neat1 specific probe from the set of A probes is used separately in RNA pull-down experiment and compared to a non-specific probe (NSP). Six Neat1 target RNAs (Canx, Fkbp4, Nono, Prl, Rbm14 and Malat1) as well as a non-target RNA (Rpa1) are analyzed by RT-qPCR. Enrichments relative to the NSP are normalized to the relative amount of corresponding Neat1_part. *p<0.05, **p<0.01, ***p<0.001.

Since the 5’ end of Neat1 appeared the main region of this lncRNA to be engaged in RNA-RNA direct interactions, we used RNA interaction prediction algorithms (IntaRNA, http://rna.informatik.uni-freiburg.de/) (24) to model the probable interactions with the 6 RNAs targets mentioned above (Figure 6 – Supplemental Table 5). Four main zones involved in the interactions with RNAs were delineated, namely Neat1-zone1 (0-190nt), Neat1-zone2 (370-420 nt), Neat1-zone3 (680-830 nt) and Neat1-zone4 (1890-2080 nt). Among these 4 zones, only Neat1-zone1 was able to bind the 6 RNAs tested. Furthermore, while Neat1-zone1 was shown to be G-rich (37%), the nucleotide composition of the 3 other zones didn’t display any specificity. In addition, none of the 4 zones contained interspersed repeat as determined by using the program RepeatMasker (http://www.repeatmasker.org). Finally, after alignment no consensus sequence was found in the 6 RNA targets that can bind any of the 4 zones of Neat1.

**Figure 6.**
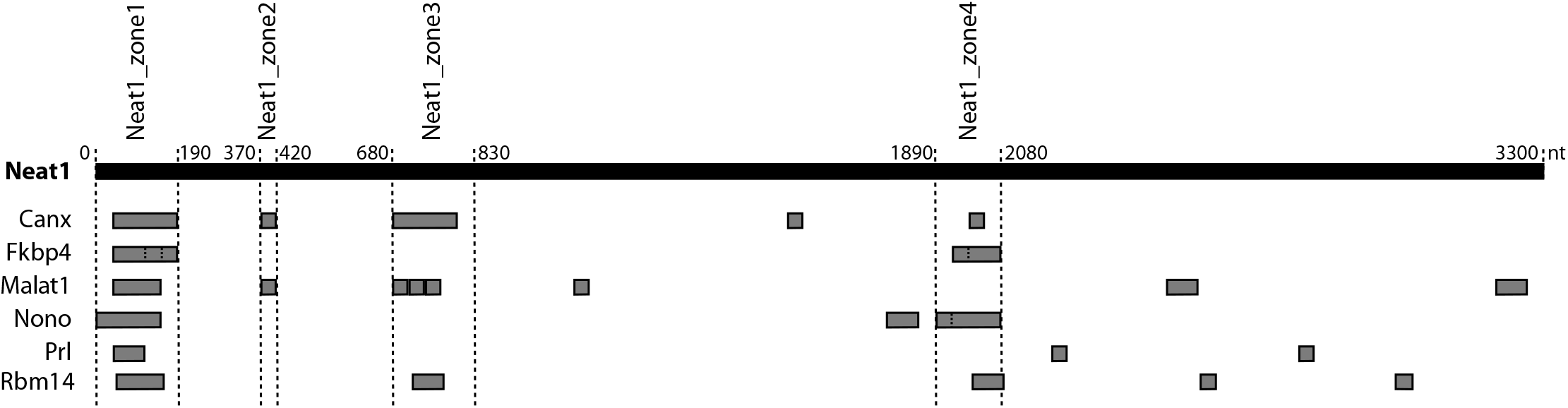
Delineation of 4 zones in the 5’ region of Neat1 involved in direct RNA interactions. The RNA interaction prediction algorithms (IntaRNA) identifies four zones of interaction between the 5’ region of Neat1 (first 3300 nucleotides) and the 6 Neat1 target RNAs (Canx, Fkbp4, Malat1, Nono, Prl and Rbm14) shown above to interact directly with the 5’ region of Neat1.

## DISCUSSION

The currently known functions of paraspeckles are to sequester not only proteins but also RNAs into the nucleus, allowing them to regulate gene expression at two different levels. By the nuclear sequestration of proteins involved in the modulation of transcription, paraspeckles are transcriptional regulators. But since paraspeckles also bind RNAs, leading to their nuclear retention that prevents their export to the cytoplasm and thus their translation into proteins (12,16), paraspeckles are also post-transcriptional regulators. The mechanism by which paraspeckles bind RNAs is however poorly understood even if two motifs in RNAs able to bind to paraspeckles have been described. The first one corresponds to an IRAlu sequence whereas the second one is a 15-nucleotide sequence, and both are located in 3’UTR of some mRNAs that are paraspeckles targets. The IRAlu sequence and the 15-nucleotide sequence bind RBPs, NONO (25) and HNRNPK (18), respectively. However, while we have previously established that in the GH4C1 cells, 4268 RNAs are targeted by paraspeckles, no IRSINE-like structure, the rat equivalent of primate IRAlus, has been found in the 3’UTR of these RNAs and only 30% of them displayed the 15-nucleotide sequence recognized by HNRNPK (18). This indicates that other mechanisms are the support of the binding of RNAs to paraspeckles.

Although it is highly likely that RNA binding occurs mostly via paraspeckle RBPs, it is also possible that the lncRNA Neat1-2 itself, which is an architectural RNA (arcRNA) necessary for the formation and maintain of paraspeckles (26), is also involved in the targeting of RNAs. It was previously demonstrated that Neat1 can sponge many different miRNAs and then can impair their functions (27). In the present study, we demonstrate that Neat1 acts not only as an arcRNA but also by the mean of direct RNA-RNA interactions, grandly contributes to the nuclear retention of RNAs and may then be considered as a post-transcriptional regulator. Indeed, the combined use of psoralen as crosslinker with RNA pull-down experiments (17) followed by RNA sequencing allowed us to identify numerous transcripts that bind directly Neat1. Confidence in the list of 1791 mRNAs directly targeted by Neat1 reported here is supported by the important overlap of targets identified by use of two different pools of Neat1-specific probes. Importantly, these direct RNA targets of Neat1 are shown to account for around 33% of the total paraspeckle targets we have previously reported in this cell line (12). It then appears that unexpectedly, no less than one third of RNAs bound to paraspeckles are engaged in direct RNA-RNA interaction with Neat1. It is also of note that in agreement with a previous study (28) we have identified by qPCR the lncRNA Malat1 (which is not officially annotated in the rat genome) as a direct RNA targets of Neat1.

Around 20% of the direct RNA targets of Neat1 are however not included in the list of total RNA targets of paraspeckles we have previously reported (12). Of course, improvement of RNA sequencing methods may account for this discrepancy. However, whereas the crosslinker reagent paraformaldehyde used in our previous study is known to allow the fixation of both the interactions between proteins and those between protein and RNA or between two RNAs caged by proteins (29), it may be suggested that it is not efficient enough to fix all RNA-RNA interactions especially those which are not in close proximity to proteins. Whatever the reason for the current identification of new RNAs as paraspeckle targets, our results allow to complete their list.

Data from the David bioinformatics resources analysis of the Neat1 RNA direct targets are consistent with the biological process and pathways in which Neat1 and paraspeckles have been shown to be involved. Therefore the enrichment in terms such as circadian rhythms (12, 30, 31), lactation (32), cell cycle in normal and cancer cells (33), and miRNAs in cancers (34) not only reinforces the link between these biological functions and paraspeckles but also provides evidence that Neat1 itself by means of direct interactions with RNAs plays a crucial role in these biological processes. While RNA splicing is more frequently associated with speckles than paraspeckles several studies have characterized a link between paraspeckles and splicing. Shut down of Neat1 has been associated with a decrease in the phosphorylation of a splicing protein (35) and up-regulation of Neat1 in lung metastases leads to a decrease in RNA splicing (36). Since among biological processes enriched in the list of direct RNA targets of Neat1, we found RNA splicing, it is tempting to speculate that some of these direct RNA targets contribute to the involvement of Neat1 in splicing mechanisms. In this view, Malat1, shown here to be a direct target of Neat1, may support the link between paraspeckles and splicing.

By designing several specific probes directed against different parts of Neat1 throughout its length and by performing Neat1 RNA pull-down experiments with a single probe, we showed that each probe is able to bind a specific part of Neat1. This result is not only, consistent with the fact that, in contrast to human Neat1-2, the rat Neat1-2 has little self-interaction (37), but it also provide evidence that despite the semi-extractability of Neat1-2 previously reported (38), our RNA-pull down protocol allows the enrichment of all parts of Neat1-2. Furthermore, being able with one probe to retrieve a single specific part of Neat1 allowed us to discriminate the different parts of Neat1-2 involved in the direct interaction with the target RNAs. Actually, the preferential location of RNA interactions corresponds to the 5’ region of Neat1-2 and possibly but to a lesser extent to the 3’ region. It is noteworthy that the central region of paraspeckles contains high concentration of proteins that constrains movements in contrast to peripheric regions, including the extremities of Neat1-2 that are more mobile (39). It should be noticed that the preferential location of RNA interactions in the 5’ region of Neat1 renders impossible to discriminate which isoform of Neat1 is involved in direct interaction with RNA targets since the 5’ region of Neat1 corresponds either to Neat1-1 or to the 5’end of Neat1-2. In any case however it is assumed that direct RNA targets are associated with the shell of paraspeckles.

Within the 5’ region of Neat1, 4 zones of interaction with different selected RNA targets have been predicted by RNA-RNA interaction prediction algorithms (IntaRNA) (24); nevertheless none of these zones displays any known feature of interaction or any retroelement according to the ReapeatMasker site. Furthermore by comparing the sequences of interaction for the selected RNA targets, we were unable to find a consensus sequence involved in the interaction with Neat1 probably because a simple motif recognition site to predict complex interaction events may be overly simplistic and provide an inaccurate view of the secondary or tridimensional structures of each RNA that are necessary for RNA-RNA interactions.

In summary, this study shows that Neat1-2 is not only involved in the formation and maintain of paraspeckles, but also contributes importantly to one of their major functions, namely to retain mRNA within the nucleus. While more than 30% of the RNA targets of paraspeckles are engaged in direct interaction with Neat1 itself, mainly through interactions occurring in the 5’ region of Neat1 and possibly but to a lesser extent in its 3’ region, it is tempting to speculate that for one same RNA, different mechanisms can be implemented to ensure its anchoring to the paraspeckles. Whether among direct RNA targets of Neat1 identified here some may also bind paraspeckle proteins to strengthen the nuclear retention remains to be determined.

## SUPPLEMENTARY DATA

Supplementary Data are available.

