## Supplemental Fig1 and Fig2 for "Direct RNA-RNA interaction between Neat1 and RNA targets, as a mechanism for RNAs paraspeckle retention"

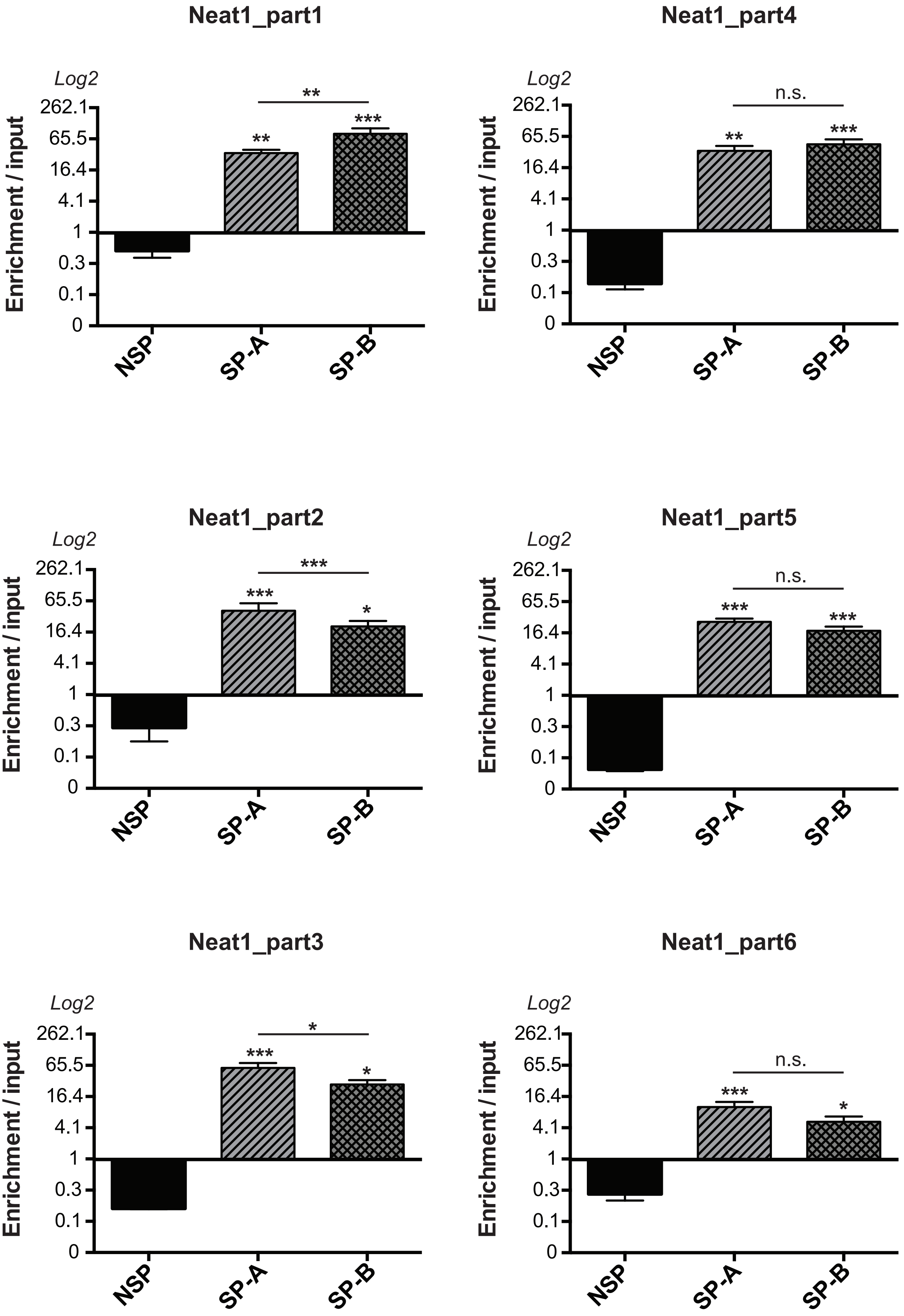
Neat1_part1
Log2
Enrichment / input
262.1
65.5
16.4
4.1
1
0.3
0.1
0
**
**
***
NSP
SP-A
SP-B
Neat1_part4
Log2
Enrichment / input
n.s.
**
***
NSP
SP-A
SP-B
Neat1_part2
Log2
Enrichment / input
***
***
*
NSP
SP-A
SP-B
Neat1_part5
Log2
Enrichment / input
n.s.
***
***
NSP
SP-A
SP-B
Neat1_part3
Log2
Enrichment / input
*
***
*
NSP
SP-A
SP-B
Neat1_part6
Log2
Enrichment / input
n.s.
***
*
NSP
SP-A
SP-B

**Supplemental Fig. 1: Pull-down of the different parts of Neat1**

Enrichment of the different parts of Neat1 (1 to 6) relative to input after RNA pull-down with a non-specific probe (NSP) or the pool of 6 probes from the set A (SP-A) or B (SP-B). Each part of Neat1 is significantly more enriched after use of SP-A or SP-B compared to NSP (part1: $F_{2,56}$ = 24,47 $p<0.0001$; part2 : $F_{2,29}$ = 38.05 $p<0.0001$ ; part3 : $F_{2,29}$ = 20.52 $p<0.0001$; part4 : $F_{2,28}$ = 16.63 $p<0.0001$ ; part5 : $F_{2,43}$ = 32. 85 $p<0.0001$ ; part6 : $F_{2,33}$ = 14.64 $p=0.0001$). Enrichment obtained is significantly different between SP-A and SP-B for Neat1_part1, Neat1_part2 and Neat1_part3. *$p<0.05$, **$p<0.01$, ***$p<0.001$vs non-specific oligonucleotide probe.

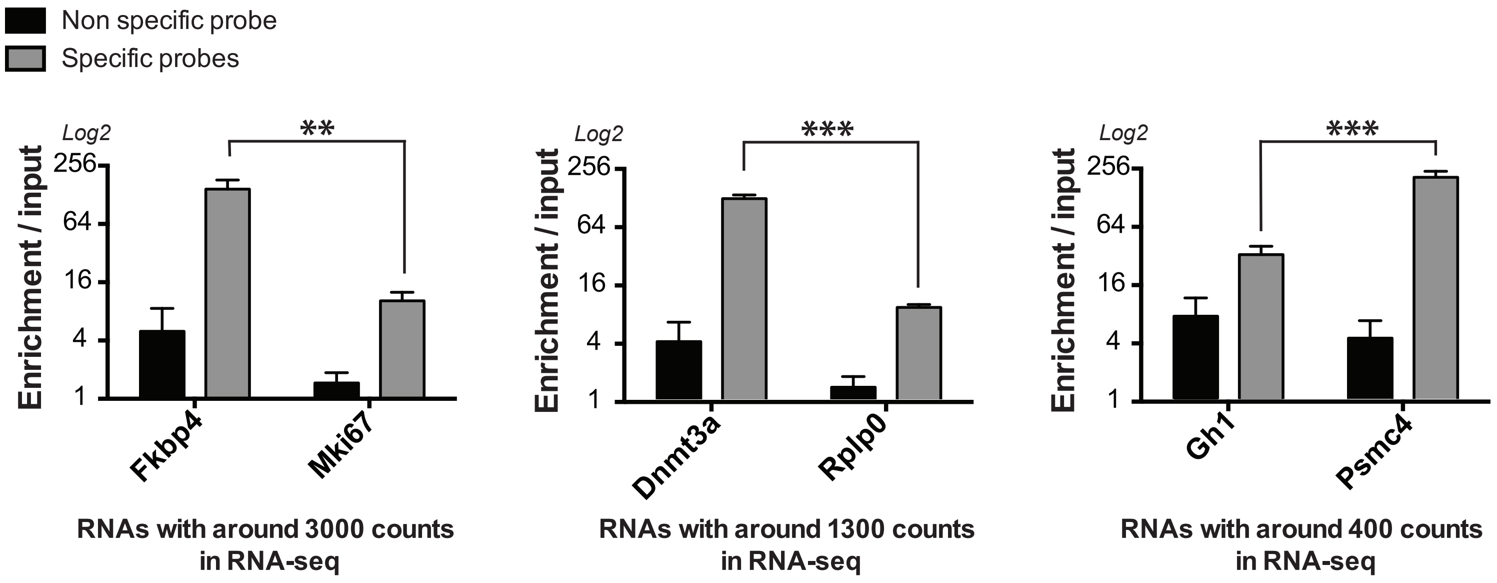
Non specific probe
Specific probes
Log2
256
64
16
4
1
Enrichment / input
**
Fkbp4
Mki67
RNAs with around 3000 counts in RNA-seq
Log2
256
64
16
4
1
Enrichment / input
***
Dnmt3a
Rplp0
RNAs with around 1300 counts in RNA-seq
Log2
256
64
16
4
1
Enrichment / input
***
Gh1
Psmc4
RNAs with around 400 counts in RNA-seq

**Supplemental Fig. 2: Magnitude of enrichment according to the number of counts in RNA-seq**

Enrichments relative to input obtained after pull-down with the pool A of specific probes (grey bar) and a non-specific probe (black bar) are compared between two RNAs displaying the same range of count number (3000, 1300 or 400 counts) (3000 counts: $F_{1,20}$=13.8, p=0.0014; 1300 counts: $F_{1,20}$=95.18, p<0.0001; 400 counts: $F_{1,20}$=29.24, p<0.0001); **p<0.01, ***p<0.001.
